# Computational modelling of atherosclerosis: developing a community resource

**DOI:** 10.1101/256750

**Authors:** Andrew Parton, Victoria McGilligan, Maurice O’Kane, Steven Watterson

**Affiliations:** Northern Ireland Centre for Stratified Medicine, Ulster University; Western Health and Social Care Trust, Altnagelvin Hospital Northern Ireland Centre for Stratified Medicine, Ulster University

**Keywords:** atherosclerosis, cardiovascular disease, computational modelling, systems biology

## Abstract

**Rationale:** Atherosclerosis is a dynamical process that emerges from the interplay between lipid metabolism, inflammation and innate immunity. The arterial location of atherosclerosis makes it logistically and ethically difficult to study *in vivo*. To improve our understanding of the disease, we must find alternative ways to investigate its progression. There is currently no computational model of atherosclerosis openly available to the research community for use in future studies and for refinement and development.

**Objective:** Here we develop the first predictive computational model to be made openly available and demonstrate its use for therapeutic hypothesis generation.

**Methods and Results:** We compiled a dataset of relevant interactions from the literature along with available parameters. These were used to build a network model describing atherosclerotic plaque development. A visual map of the network model was produced using the Systems Biology Graphical Notation (SBGN) and a dynamic mathematical description of the network model that enables us to simulate plaque growth was developed and is made available using the Systems Biology Markup Language (SBML). We used this model to investigate whether multi-drug therapeutic interventions could be identified that stimulate plaque regression. The model produced comprised 20 cell types and 41 proteins with 89 species in total. The visual map is available for reuse and refinement using the SBGN Markup Language standard format and the mathematical model is available using the SBML standard format. We used a genetic algorithm to identify a multi-drug intervention hypothesis comprising five drugs that comprehensively reverse plaque growth within the model.

**Conclusions:** We have produced the first predictive mathematical and computational model of atherosclerosis that can be reused and refined by the cardiovascular research community. We demonstrated its potential as a tool for future studies of cardiovascular disease by using it to identify multi-drug intervention hypotheses.

**Subject Codes:** Atherosclerosis, Computational Biology, Lipids and Cholesterol, Cell Signaling/Signal Transduction, Cardiovascular Disease

## 1. Introduction

Cardiovascular disease (CVD) is the primary cause of global mortality. CVD is estimated to account for 17m deaths worldwide each year, representing 31% of all cause mortality worldwide and 47% of all cause mortality within Europe^a^. Such a prevalent condition incurs a significant financial burden, accounting for 17% of all healthcare expenditure in the USA^1^. Age is a significant risk factor and with an aging population, the cost of CVD related therapies is predicted to almost triple in the USA from $273 billion in 2010 to $818 billion by 2030^1^.

Atherosclerosis is estimated to account for 71% of CVD diagnoses^a^. It is characterised by the hardening of an artery wall, and the formation of a fibrous-fatty lesion within the intimal layer. As the disorder progresses, thick extracellular cores of lipid build within the artery wall, occluding the artery and subsequently reducing blood flow. Thrombosis can further occlude the artery either as a result of plaque rupture or turbulent blood flow induced around the site of the atheroma.

Despite our increasing knowledge of the mechanisms driving this disorder, the pathogenesis of atherosclerosis is still not fully understood. In part, this is due to the significant challenge inherent in studying live, dynamic plaques. Accessing plaques *in vivo* is logistically difficult, necessitating catheterization, and ethically challenging as it can increase the risk of plaque rupture. As a result, alternative approaches to studying atherosclerosis dynamics are needed. Computational modelling has the potential to be especially valuable here due to its flexibility, low financial and ethical cost, consistency and ease of replication. However, currently there are no computational or mathematical models of atherosclerosis that are easily available to the research community for use in exploratory studies.

In previous modelling studies the majority of work has focused on plaque initiation and haemodynamics^2^, where Navier-Stokes dynamics have described blood flow and wall shear stress has been calculated as an pro-atherogenic output^3^. We are interested in the molecular and cellular biology that mediate plaque formation and can furnish targets for therapeutic interventions. However, in previous studies these details have been routinely omitted or simplified for reasons of mathematical expediency^4^. Furthermore, the resulting models have not been made publicly available. Reusing this work would necessitate reconstruction of the models in their entirety, a complex, time consuming and error-prone task. At the present time, the European Bioinformatics Institute (EBI) Biomodels database^b^ contains only one model pertaining to atheroma formation, focussing on lipoprotein action and B-cell signaling with little detail on the mechanisms of plaque formation^5^. KEGG^6^, Reactome^7^ and Wikipathways^8^ contain no molecular biology maps of atherosclerosis. Here we develop the first detailed, predictive dynamical computational model of atherogenesis using Systems Biology standards. The model comprises a map composed using the Systems Biology Graphical Notation (SBGN)^9^ and made available to the research community for reuse and refinement using the Systems Biology Graphical Notation Markup Language (SBGN-ML)^10^. This map is accompanied by a mathematical model describing the dynamics of the interactions in the map as a system of ordinary differential equations (ODEs), and made available using the Systems Biology Markup Language (SBML)^11^. There are many examples of SBGN^c^ and SBML^d^ compliant software.

Currently, treatment of atherosclerotic vascular disease focuses on limiting disease progression (though smoking cessation, lipid lowering and anti-platelet therapies and optimal management of hypertension and diabetes) and revascularistion procedures such as angioplasty and bypass grafting to clinically relevant stenotic lesions in the coronary, peripheral or cerebral vasculature. Although such treatments are clinically effective, it is less clear whether medical therapies can reduce plaque size, although there is some evidence to suggest that intense statin treatment^12^, combined statin-PCSK9 inhibitor treatment^13^ or Cyclodextrin treatment^14^ may yield a modest plaque reduction. New drug combinations that yield a substantial reduction in plaque size could have a dramatic impact on CVD morbidity and mortality and so their identification has high strategic importance. Here, we employ the model to develop effective therapeutic hypotheses comprising multi-drug combinations.

## 2. Methods

A list of the cell types involved in atherosclerosis was compiled from the existing literature (see supplementary table 4). Each article identified was also searched for references to proteins and small molecules with each entity found considered for the model. A protein or small molecule was incorporated into the model if its biological source, presence within a relevant compartment and its influence on atherogenesis (however minor) were all described. The model was assembled with CellDesigner^15^ using the SBGN schema and with mass action and Michaelis-Menten equations primarily used to describe the dynamics. The resulting model was exported to SBGN-ML file format to disseminate the visual map and to SBML file format to disseminate the mathematical model describing the dynamics.

PubMed and Google Scholar searches were undertaken to find studies describing representative concentrations of the cells, proteins and small molecules. The BRENDA enzyme database was searched for relevant known rate parameters^16^. Values for unknown parameters were calculated by constraining the model to show dynamics in agreement with published CVD studies. We considered dynamics for three lipid profiles: high risk, medium-risk and low-risk comprising LDL concentrations of 190 mg/dl^e^, 110 mg/dl^e^ and 50mg/dl^17^, respectively and HDL concentrations of 40 mg/dl, 50 mg/dl and 50 mg/dl, respectively^18^. Atherosclerosis is considered to be a chronic condition. Hence, we considered plaque formation across a representative time scale of 80 years.

There are between 5 and 800 cells within a plaque area per high powered field (HPF) at 400× magnification^19^, where one HPF displays approximately 0.2mm^2^ of plaque area^20^. We estimate that a plaque contains between 25 and 4000 cells per mm^2^. Average plaque area has been shown to be 15.2mm^2^ (^21^), giving the number of cells in a plaque as being between 380 and 60800. With this, we identified the following constraints from the published literature.

I) Smooth muscle cells comprise 35.10% of the cellular composition of plaques^20^, corresponding to a range of 133 cells which we take to be representative of low LDL profiles to 21341 cells which we take to be representative of high LDL profiles.

II) Macrophages (including foam cells) comprise 34.07% of the cellular composition of plaques^20^, corresponding to a range of 129 cells to 20715 cells.

III) The ratio of Th1 to non-Th1 cells in a plaque is approximately 0.3^22^, corresponding to a range of 88 Th1 cells to 14031 Th1 cells.

IV) Blood serum concentrations of MCP1/CCL2 were estimated from myocardial infarction and ischemic stroke patients, ranging from 100 pg/ml to 775 pg/ml^23^.

V) Blood serum concentrations of CXCL9 were estimated from patients assessed for coronary artery calcium deposits, ranging from 17.4 pg/ml to 271.2 pg/ml^24^.

VI) Blood serum concentrations of CXCL10 were estimated from patients assessed for coronary artery disease, ranging from 127.6 pg/ml to 956.5 pg/ml^25^.

VII) Blood serum concentrations of CXCL11 were estimated from control groups in transplantation studies, ranging from 420 pg/ml to 1062 pg/ml^26^.

VIII) Blood serum concentrations of I L1b were estimated from congestive heartfailure and control patients, ranging from 0.28 pg/ml to 2.12 pg/ml^27^.

IX) Plaque concentrations of TIMP1 were estimated from carotid endarterectomy patients, ranging from 5.3 μg/g to 12.4 μg/g wet weight of plaque^28^.

X) Plaque concentrations of IFNg were estimated from carotid endarterectomy patients, ranging from 20 pg/g to 182 pg/g wet weight of plaque^29^.

XI) Plaque concentrations of TGFb were estimated from control and coronary artery disease patients, ranging from 0.33 mg/g to 0.76 mg/g of protein^30^.

XII) Plaque density ratios of chymase to tryptase were recorded to be 107.8:135.1 in plaques^31^.

XIII) T Cells comprise 30.82% of the cellular composition of plaques^20^, corresponding to a range of 117 cells to 18739 cells.

XIV) Blood serum concentrations of CCL5 were estimated from control and coronary event patients, ranging from 2.7 ng/ml to 176.0 ng/ml, respectively^32^.

XV) Plaque concentrations of MMP1 were estimated from carotid endarterectomy patients, ranging from 18 ng/g to 104 ng/g wet weight of plaque^28^.

XVI) Plaque concentrations of MMP9 were estimated from carotid endarterectomy patients, ranging from 121 ng/g to 722 ng/g wet weight of plaque^28^.

XVII) Plaque concentrations of IL1b were estimated from carotid endarterectomy patients, ranging from 12 ng/g to 24 ng/g wet weight of plaque^28^.

XVIII) Plaque concentrations of IL6 were estimated from carotid endarterectomy patients, ranging from 1.5 μg/g to 5.1 μg/g wet weight of plaque^28^.

XIX) Plaque concentrations of TNFa were estimated from carotid endarterectomy patients, ranging from 15 ng/g to 27 ng/g wet weight of plaque^28^.

XX) Plaque concentrations of IL10 were estimated from arterial occlusion patients, ranging from 1.51 pg/mg to 2.29 pg/mg wet weight of plaque^33^.

XXI) Plaque concentrations of IL12 were estimated from arterial occlusion patients, ranging from 3.6 pg/mg to 4.6 pg/mg wet weight of plaque^33^.

XXII) Plaque concentrations of elastin were estimated from acute coronary syndrome patients, giving 1.58 mg/g wet weight of plaque^34^.

XXIII) Plaque concentrations of collagen were estimated from acute coronary syndrome patients, giving 6.26 mg/g wet weight of plaque^34^.

XXIV) Plaque concentrations of PDGF were estimated from carotid endarterectomy patients, ranging from 279 pg/g to 1381 pg/g wet weight of plaque^29^.

XXV) The weight of oxidized LDL per weight of ApoB has been measured to be 19.6 ng/μg in macrophage rich plaques and 1.9 ng/μg in normal intimal tissue^35^. The plaque concentration of ApoB has been measured to range from 1.97 μg/mg to 0.13 μg/mg^36^, yielding upper and lower estimates for oxidised LDL concentrations of 38.6 μg/g and 0.25 μg/g.

XXVI) Plaque concentrations of IL2 were estimated from acute coronary syndrome patients, giving 24.0 pg/mg of protein^37^.

XXVII) Plaque concentrations of IL18 were estimated from acute coronary syndrome patients, giving 10.7 pg/mg of protein^37^.

XXVIII) Blood serum concentrations of chylomicrons were estimated from a control group and hyperlipidemic patients, corresponding to 1.4 μg/ml and 52.6 μg/ml, respectively^38^.

XXIX) Blood serum concentrations of triglycerides were estimated from a control group and hyperlipidemic patients, corresponding to 58 mg/dl and 1005 mg/dl, respectively^38^.

XXX) The ratio of Th1 to Th2 cells has been shown to correlate with atherogenesis^39^.

XXXI) Animal models with advanced atherosclerosis have shown plaque reduction mediated by reverse cholesterol after a reduction in lipid profile^40^.

After an initial model was constructed with the known parameter values, the unknown parameters were optimised to ensure that the model adhered to these experimental results as far as possible.

### Multi-drug plaque regression therapeutic hypotheses

In order to demonstrate the utility of the model, we undertook to identify an optimal multi-drug intervention hypothesis that would reprogram the dynamics of the model leading to regression of advanced plaques. It has been demonstrated that multidrug approaches have the potential to exploit compound effects to yield effective interventions at lower individual and collective dosages than in comparable single-drug interventions, reducing the risk from pleotropic effects^41^. This is an example of the type of investigation that would be extremely complex to undertake clinically and yet can be undertaken computationally with relative ease.

We identified the following 9 drugs with targets in the model (proteins they inhibit in brackets): 2-(4-Chloro-3-(trifluoromethyl)phenoxy)-5-(((1-methyl-6-morpholino-2-oxo-1,2-dihydropyrimidin-4-yl)oxy)methyl)benzonitrile (PLA2), GW4869 (SMase), Quercetin Monoglucoside (Lipoxygenase), cFMS Receptor Inhibitor III (MCSF), Bindarit (CCL2), Imatinib Mesylate (PDGF), Ustekinumab (IL12R), GSK1070806 (IL18R), SCH546738 (CXCL9, CXCL10, CXCL11, CCL5). Concentrations were presented as multiples of the corresponding k_i_ and because PLA2, SMase and Lipoxygenase all catalyse the same interaction, we constrained these drugs to have the same concentration, giving a set of drugs with seven degrees of freedom.

We used the MATLAB^f^ software system and a genetic algorithm with a population size of 10000 for 100 generations to identify the optimal combination of drugs that would drive atherosclerosis regression. The genetic algorithm started from one instance of a set of drug concentrations and from this generated a further 9999 instances of sets of drug concentrations from the first by adding Gaussian noise to the concentration of each drug (with standard deviation 1, the default setting). These 10000 instances comprised the first generation of candidate interventions. All instances were evaluated for their efficacy at plaque reduction and 10000 new instances were created as a second generation of candidate interventions from the two most effective instances of the first generation (with the addition of Gaussian noise). The 10000 new instances were then themselves evaluated with the two most effective instances being used to generate a further 10000 new instances, the third generation. This process was iterated until we arrived at instances from which no improvement in efficacy could be found at which point the best performing instance was interpretted as optimal. In order to evaluate the efficacy of a particular instance, we constructed a scoring function that allowed the model to develop using the high risk profile for the first forty years before introduction of the drug concentrations of the instance at forty years. The model then continued to run for a further forty years and at eighty years we calculated a score for the instance as S = (C/C_max_ + T/T_max_)/2 + 0.01*(sum of drug concentrations) where C is the sum of smooth muscle cells, macrophages, foam cells and T-cells observed and C_max_ is the sum of smooth muscle cells, macrophages, foam cells and T-cells that would occur at eighty years in the absence of any drugs. T is the collagen concentration observed and T_max_ is the collagen concentration that would occur at eighty years in the absence of any drugs. This score describes the efficacy of the instance of a set of drugs at driving plaque regression with effective interventions yielding lower numbers and ineffective interventions yielding higher numbers. The score also included the sum of the concentrations of the drugs used. Low scores also ensure that the dosages are minimal, yielding therapeutic hypotheses with reduced risks of off-target effects. At each generation, the genetic algorithm selected the two instances with the lowest scores. The analysis was run on an Intel(R) Xeon(R) CPU E5-2630 v3 @ 2.6GHz (Octo-core) CPU with 64GB of RAM running CentOS 7.

## 3. Results

A visual map of the model obtained is shown in Figure 1 using the SBGN schema. The model covers five distinct organs and tissues: the liver, intestine, lumen, endothelium and tunica intima. It covers LDL retention, LDL oxidation, monocyte recruitment, monocyte differentiation, smooth muscle cell proliferation, phagocytosis, reverse cholesterol transport and T-cell proliferation. The cell types involved include monocytes, endothelial cells, T-cells, macrophages, foam cells, B-cells, smooth muscle cells, neutrophils, dendritic cells and mast cells. A legend describing the glyphs of the SBGN schema is shown in Figure 2. Each interaction represents a parameterized equation (see supplementary table 1 for the equations and supplementary table 2 for the parameters), enabling us to dynamically simulate the changing concentrations/abundances of the model as the plaque forms.

**Figure 1.**
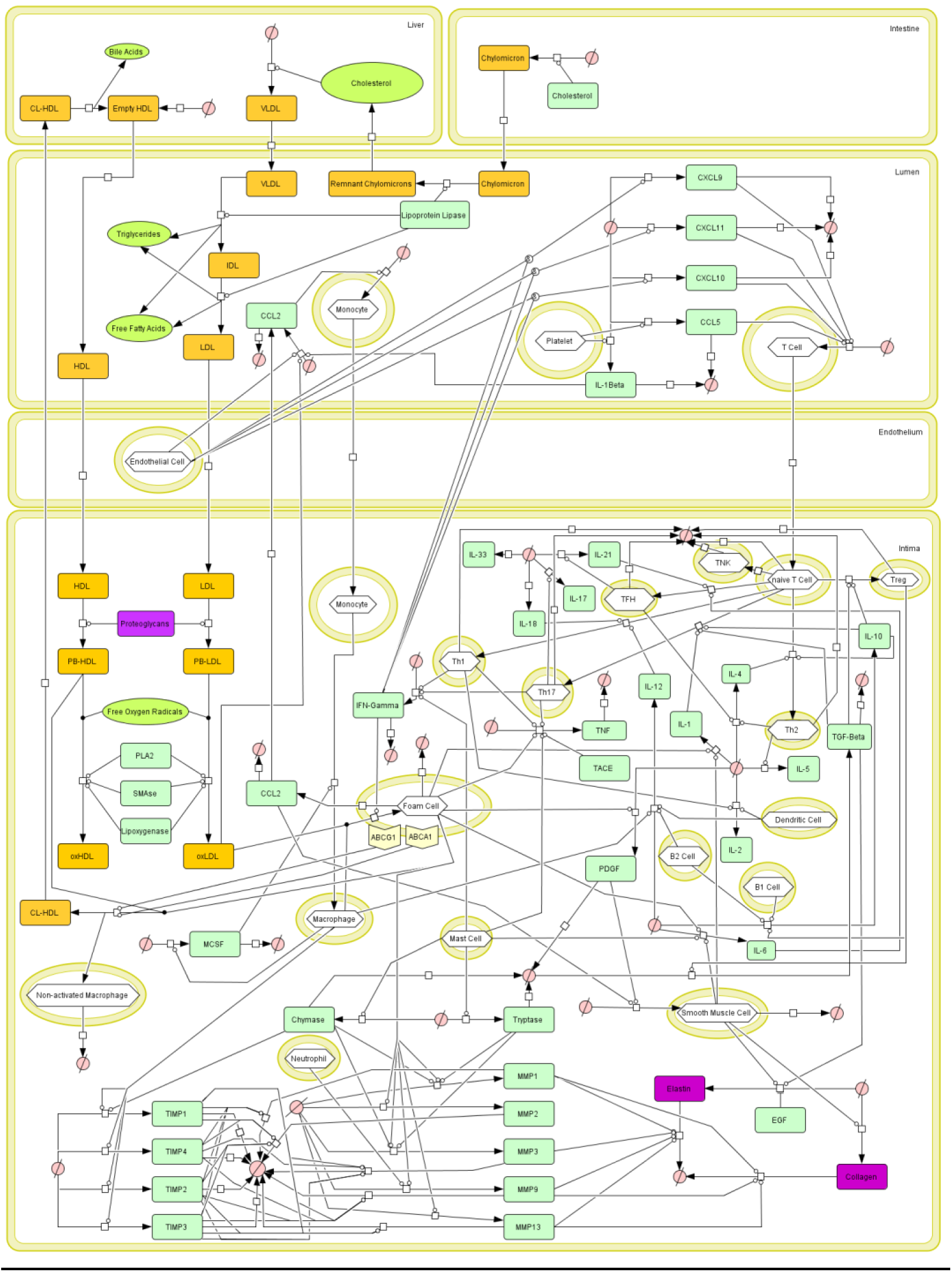
A map of atherosclerotic plaque dynamics shown using the Systems Biology graphical Notation (SBGN).

**Figure 2.**
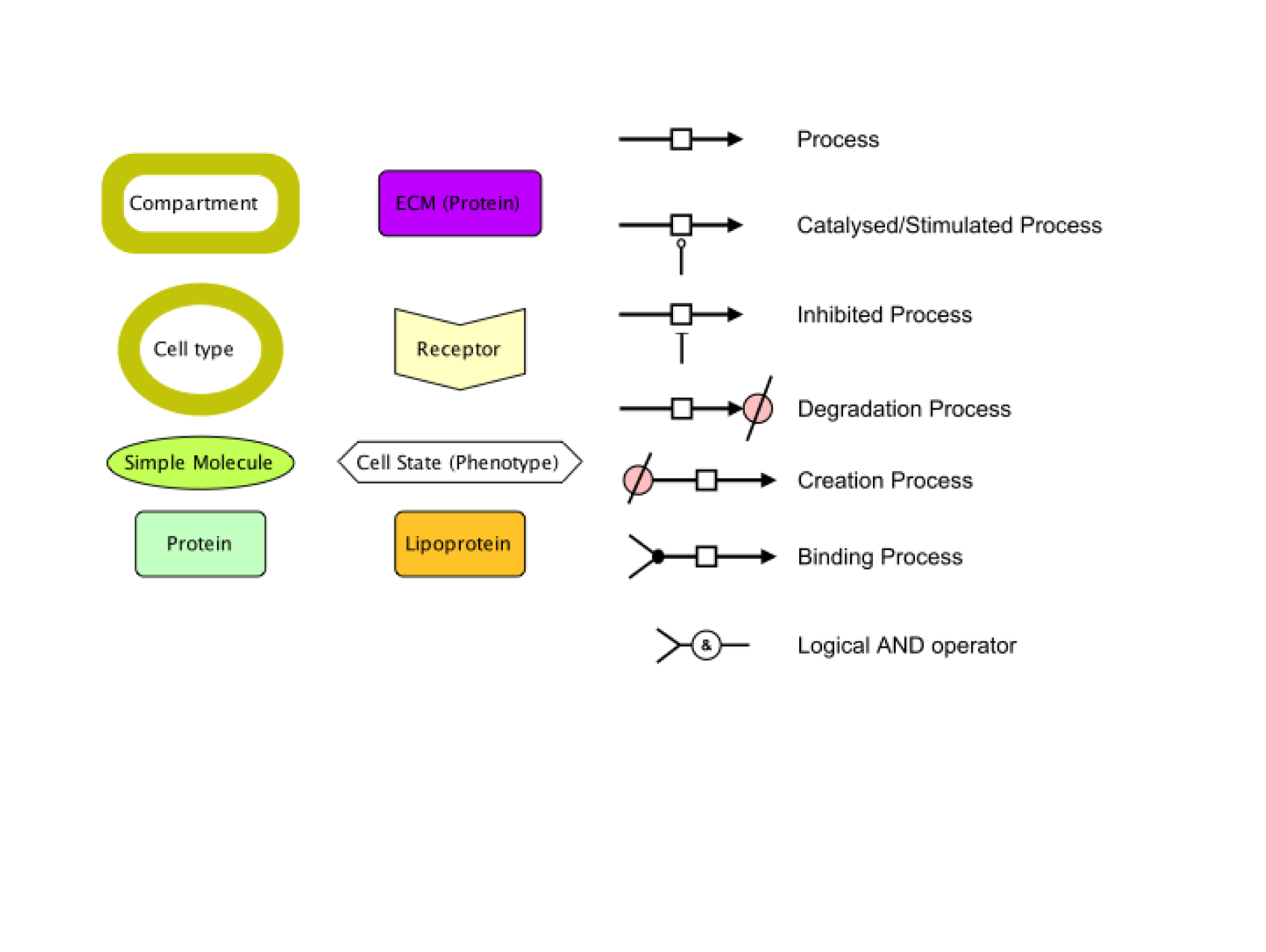
The legend for the SBGN schema used in Figure 1.

The initial conditions identified are described in supplementary table 1 and unknown parameters were optimised so that the model maximally satisfied the constraints described above simultaneously. Key markers for plaque development include smooth muscle cell, macrophage and foam cell and Th1 cell proliferation. Their behavior is shown in Figure 3 for the three risk profiles.

**Figure 3.**
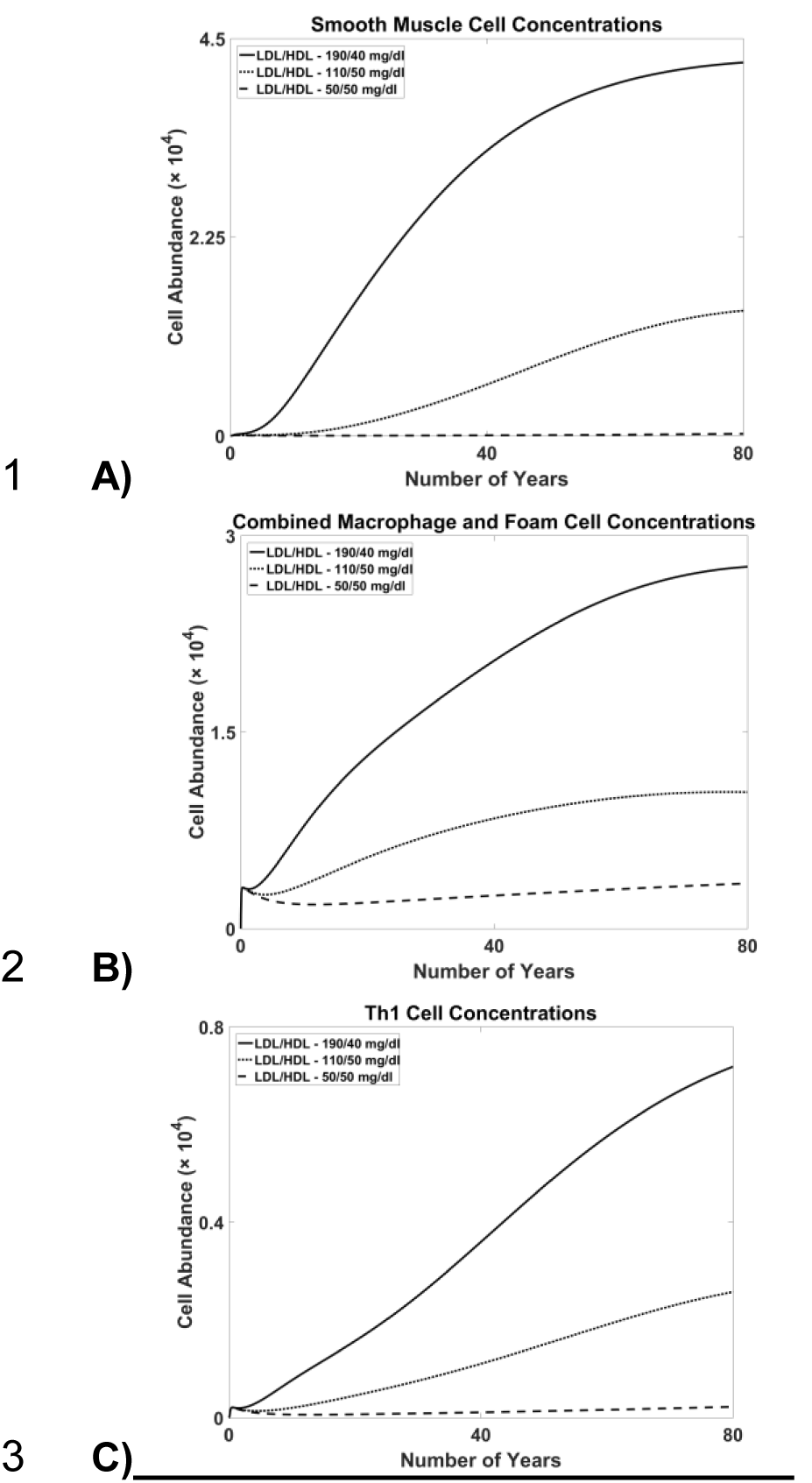
(A) Smooth muscle cell concentrations, (B) macrophage and foam cell concentrations and (C) Th1 cell concentrations during plaque development for the three blood LDL/HDL profiles 190/40 mg/dl, 110/50 mg/dl and 50/50 mg/dl.

The model satisfies the constraints as follows. Results are stated at 80 years with constraint values in brackets.

I) Figure 3A shows smooth muscle cell abundance, yielding 42287 cells (21341) and230 cells (133), for high and low risk profiles, respectively.

II) Figure 3B shows combined macrophage and foam cell abundance, yielding 27630 cells (20715) and 3463 cells (129) for high and low risk profiles, respectively.

III) Figure 3C shows Th1 cell abundance, yielding 7186 cells (14031) and 223 cells (88) for high and low risk profiles, respectively.

IV) Figure 4.1 shows MCP1/CCL2 blood serum concentration, yielding 649.8 pg/ml (775 pg/ml) and 163.8 pg/ml (100 pg/ml) for high and low risk profiles, respectively.

**Figure 4.**
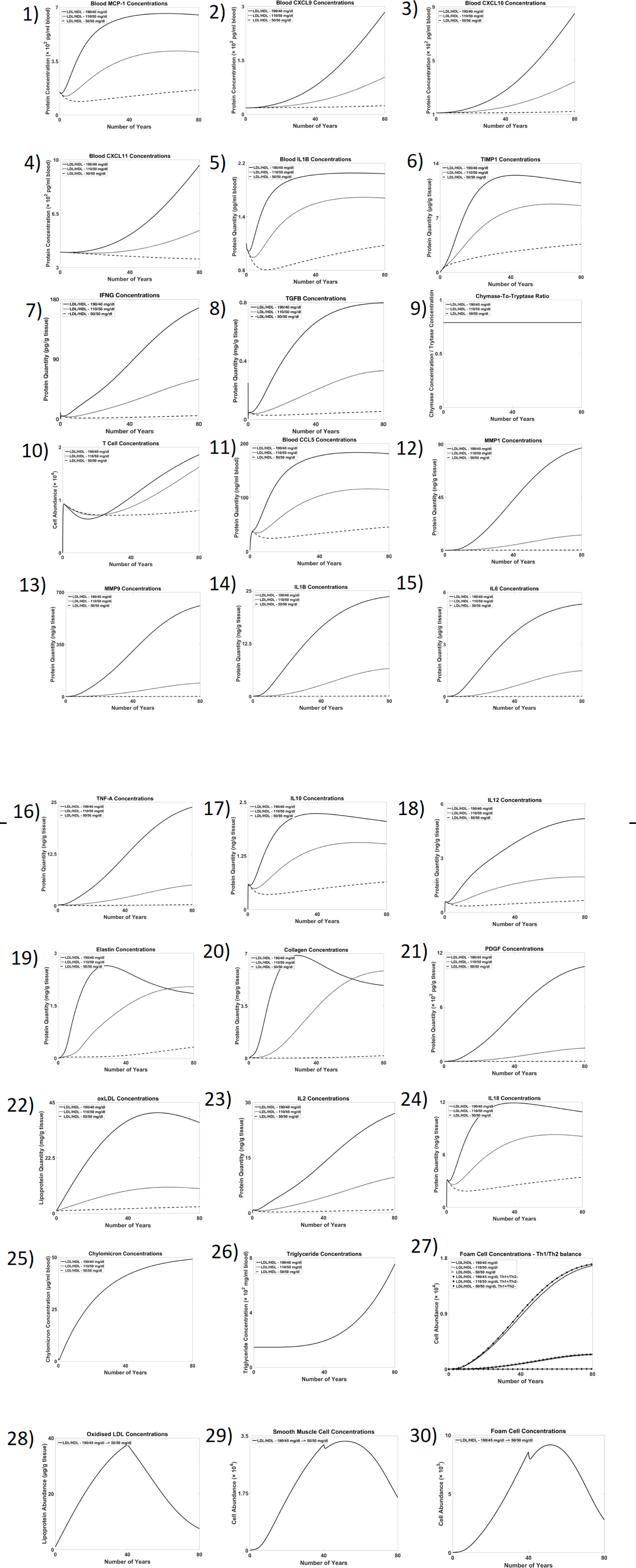
The performance of the model for clinical requirements determined from the literature.

V) Figure 4.2 shows CXCL9 blood serum concentration, yielding 283.9 pg/ml (271.2 pg/ml) and 23.8 pg/ml (17.4 pg/ml) for high and low risk profiles, respectively.

VI) Figure 4.3 shows CXCL10 blood serum concentration, yielding 850.0 pg/ml (956.5 pg/ml) and 120.9 pg/ml (127.6 pg/ml) for high and low risk profiles, respectively.

VII) Figure 4.4 shows CXCL11 blood serum concentration, yielding 965 pg/ml (1062pg/ml) and 355 pg/ml (420 pg/ml) for high and low risk profiles, respectively.

VIII) Figure 4.5 shows IL1b blood serum concentration, yielding 2.04 pg/ml (2.12pg/ml) and 0.97 pg/ml (0.28 pg/ml) for high and low risk profiles, respectively.

IX) Figure 4.6 shows TIMP1 plaque concentration, yielding 11.5 μg/g (12.4 μg/g) and 3.6 μg/g (5.3 μg/g) for high and low risk profiles, respectively.

X) Figure 4.7 shows IFNg plaque concentration, yielding 167 pg/g (182 pg/g) and 5 pg/g (20 pg/g) for high and low risk profiles, respectively.

XI) Figure 4.8 shows TGFb plaque concentration, yielding 0.80 mg/g (0.76 mg/g) and 0.05 mg/g (0.33 mg/g) for high and low risk profiles, respectively.

XII) Figure 4.9 shows the ratio of plaque density between chymase and tryptase, yielding 106.0:134.3 (107.8:135.1) for the high risk profile.

XIII) Figure 4.10 shows total T cell abundance, yielding 18562 cells (18739) and 8012 cells (117) for high and low risk profiles, respectively.

XIV) Figure 4.11 shows CCL5 blood serum concentration, yielding 181.1 ng/ml (176.0 ng/ml) and 45.7 ng/ml (2.7 ng/ml) for high and low risk profiles, respectively.

XV) Figure 4.12 shows MMP1 plaque concentration, yielding 86.8ng/g (104 ng/g) and 0.2 ng/g (18 ng/g) for high and low risk profiles, respectively.

XVI) Figure 4.13 shows MMP9 plaque concentration, yielding 609.6 ng/g (722 ng/g) and 1.6 ng/g (121 ng/g) for high and low risk profiles, respectively.

XVII) Figure 4.14 shows IL1b plaque concentration, yielding 23.6 ng/g (24 ng/g) and 0.1 ng/g (12ng/g) for high and low risk profiles, respectively.

XVIII) Figure 4.15 shows IL6 plaque concentration, yielding 5.3 μg/g (5.1 μg/g) and0.025 μg/g (1.5 μg/g) for high and low risk profiles, respectively.

XIX) Figure 4.16 shows TNFa plaque concentration, yielding 24 ng/g (27 ng/g) and 0.3 ng/g (15 ng/g) for high and low risk profiles, respectively.

XX) Figure 4.17 shows IL10 plaque concentration, yielding 2.1 ng/g (2.3 ng/g) and 0.6 ng/g (1.5 ng/g) for high and low risk profiles, respectively.

XXI) Figure 4.18 shows IL12 plaque concentration, yielding 5.2 ng/g (4.6 ng/g) and 0.7 ng/g (3.6 ng/g) for high and low risk profiles, respectively.

XXII) Figure 4.19 shows the elastin plaque concentration, yielding 1.85 mg/g (1.58mg/g) for the high risk profile.

XXIII) Figure 4.20 shows collagen plaque concentration, yielding 4.87 mg/g (6.26 mg/g) for the high risk profile.

XXIV) Figure 4.21 shows PDGF plaque concentration, yielding 1048 pg/g (1381 pg/g) and 2 pg/g (279 pg/g) for high and low risk profiles, respectively.

XXV) Figure 4.22 shows oxidised LDL plaque concentration depending on risk profile. At 80 years, the high risk profile yields 36.8 μg/g (38.6 μg/g) and the low risk profile yields 2.6 μg/g (0.25 μg/g).

XXVI) Figure 4.23 shows IL2 plaque concentration, yielding 27 ng/g (24 ng/g) for the high risk profile.

XXVII) Figure 4.24 shows IL18 plaque concentration, yielding 10.9 ng/g (10.7 ng/g) for the high risk profile.

XXVIII) Figure 4.25 shows chylomicron blood serum concentration, yielding 49.1 μg/ml (52.6 μg/ml) a value that does not change for low risk profiles (1.4 μg/ml).

XXIX) Figure 4.26 shows triglyceride blood serum concentration, yielding 754 mg/dl (1005 mg/dl) a value that does not change for low risk profiles (58 mg/dl).

XXX) Figure 4.27 shows foam cell aggregation after the parameter determining rate of differentiation to Th1 cells has been increased by 10% and the parameter determining the rate of differentiation to Th2 cells has been decreased by 10%. This has lead to a modest increase in foam cell concentrations for a high risk profile.

XXXI) Figures 4.28, 4.29 and 4.30 shows oxidized LDL concentration, smooth muscle cell and foam cell abundance, respectively, when the LDL and HDL are switched from 190 mg/dl and 40 mg/dl, respectively, to 50mg/dl and 50mg/dl, respectively, after 40 years, demonstrating plaque reduction.

In addition to addressing these constraints, the model also agrees with the following clinical results.

XXXII) Blockade of endogenous IL-12 has been shown to reduce atherogenesis^42^. Figure 5.1 shows that with a 75% reduction to the rate parameters describing IL-12 production, foam cell abundance is significantly reduced for high and mid risk profiles.

**Figure 5.**
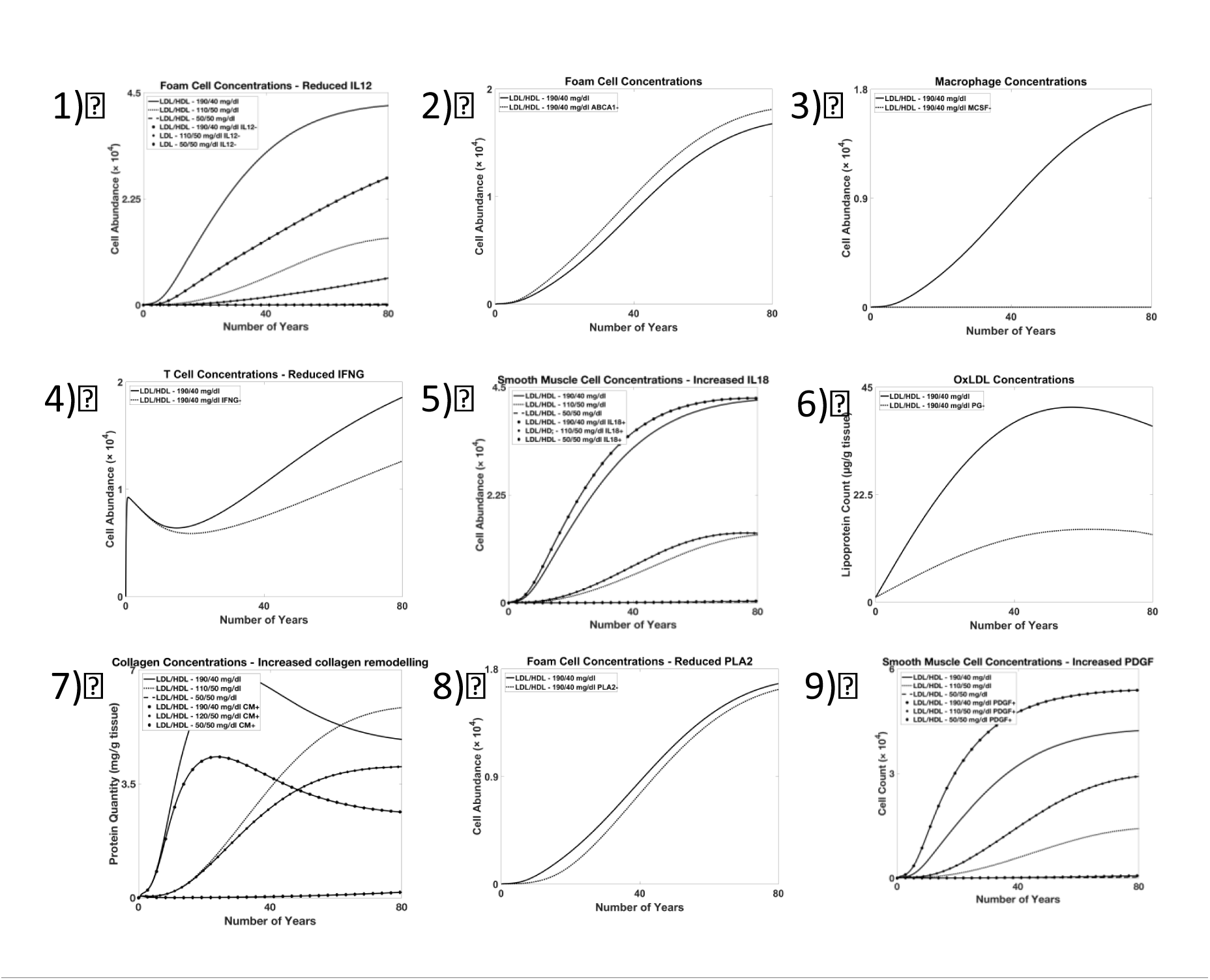
The performance of the model for further clinical observations.

XXXIII) Deficiency of ABCA1 function impairs reverse cholesterol transport and increases atheroma size^43^. Figure 5.2 shows that with a reduction in the initial

ABCA1 concentration by 90%, foam cell concentration is increased across the lifetime of the simulation.

XXXIV) Deficiency of MCSF reduces monocyte/macrophage circulation and plaqueformation^44^. Figure 5.3 shows that with a reduction in the initial MCSF concentrations from 100 μg/mg of tissue to 0, macrophage abundance drops significantly within the plaque.

XXXV) T-cells abundance is reduced as a result of IFNGR knockout^45^. Figure 5.4 shows that decreasing the k_cat_ rate parameter describing IFNG production by 50%reduces T-cell abundance within the plaque.

XXXVI) IL-18 has been shown to be atherogenic^46^. Figure 5.5 shows that increasing the rate parameter describing IL-18 production by 50%, increases smooth muscle cell recruitment within the plaque.

XXXVII) Reduction in proteoglycan concentration reduces intimal oxLDL concentrations^47^. Figure 5.6 shows that decreasing the initial concentration of proteoglycan concentration from 500 μg/mg of tissue to 100 pg/mg of tissue reduces the concentration of oxidized LDL within the plaque.

XXXVIII) Increasing activity of matrix metalloproteinases leads to degradedextracellular matrix^48^. Figure 5.7 shows that doubling the binding rate parameter between extra cellular matrix and matrix metalloproteinases significantly reduces collagen concentrations.

XXXIX) PLA2 concentration has been shown to correlate with atherogenesis^49^. Figure 5.8 shows that a reduction in the initial PLA2 concentrations by 90% reduces the foam cell concentration within the plaque.

XL) Increasing PDGF activity increases smooth muscle cell abundance^50^. Figure 5.9 shows that increasing the rate parameter describing PGDF production by 200% increases smooth muscle cell recruitment in the plaque.

### Reusability of the model

The visual map is available using the SBGN-ML file format and the mathematical model is available using the SBML file format from the supplementary material. The mathematical model is also available from the European Bioinformatics Institute–s Biomodels repository (MODEL1710020000).

The files can be opened, edited and analysed in software supporting the SBGN-ML and SBML standards. SBML compliant software includes Copasi^g^, Cytoscape with the cy3SBML plugin^h^ and Dizzy^i^. Figure 6 shows the graphical map opened in three representative SBGN compliant editors: New^t^, PathVisio^k^ and VANTED with SBGN-ED extension^l^ along with a subsection of the plain text, XML file.

**Figure 6:**
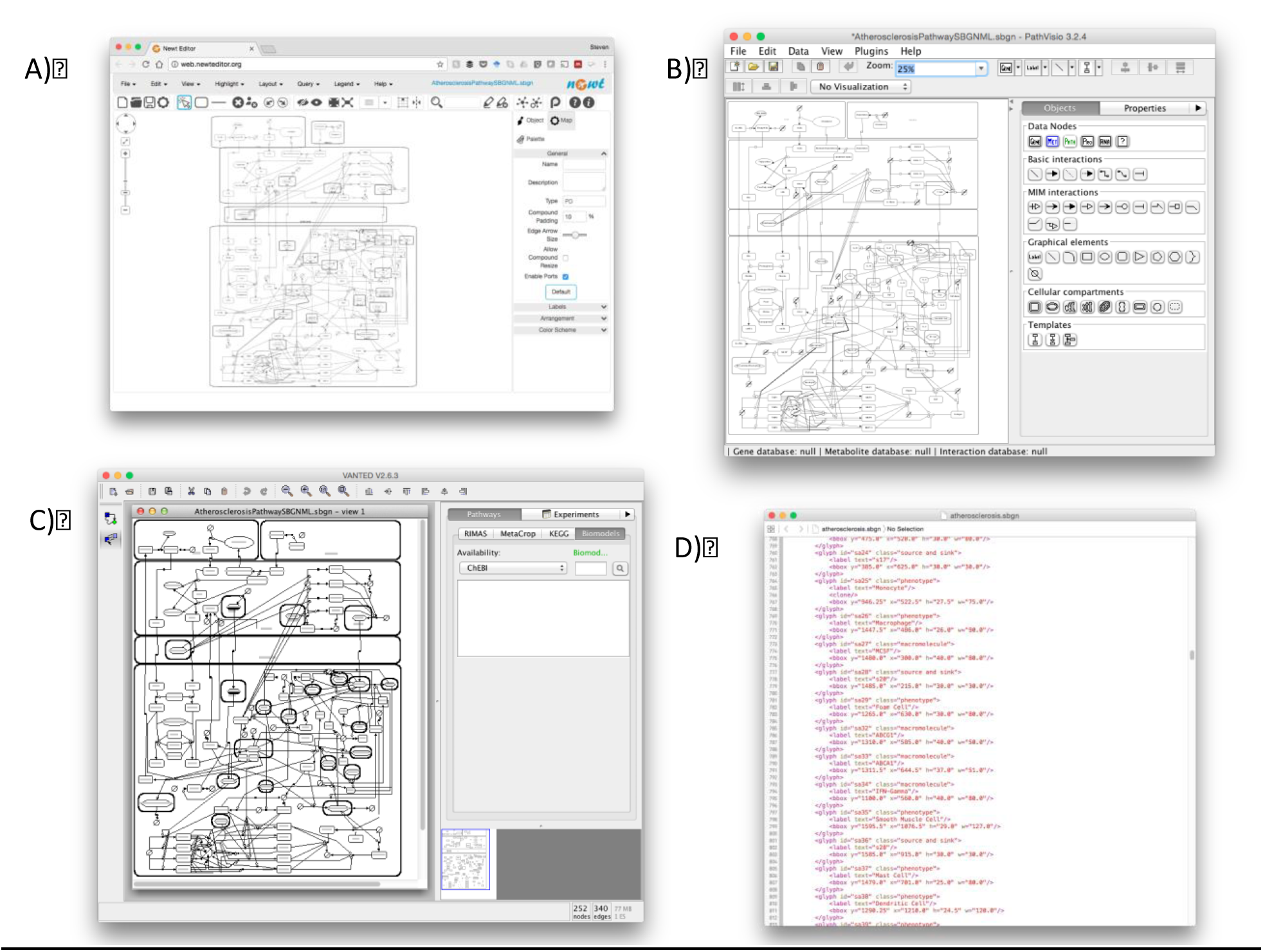
The model viewed in using the A) Newt B) PathVisio and C) VANTED platforms and D) viewed as plain text XML.

### Therapeutic hypothesis generation

We determined the following drug combination that optimally drove plaque regression: 2-(4-Chloro-3-(trifluoromethyl)phenoxy)-5-(((1-methyl-6-morpholino-2-oxo-1,2-dihydropyrimidin-4-yl)oxy)methyl)benzonitrile (PLA2) – 4.35×10^−5^, GW4869 (SMase) – 4.35×10^−5^, Quercetin Monoglucoside (Lipoxygenase) – 4.35×10^−5^, Bindarit (CCL2) – 37.0, cFMS Receptor Inhibitor III (MCSF) – 0, SCH546738 (CXCL9, CXCL10, CXCL11, CCL5) – 8.45x10^−4^, Ustekinumab (IL12R) – 7.62, GSK1070806 (IL18R) – 7.60, Imatinib Mesylate (PDGF) – 0, where, concentrations are described as multiples of the corresponding inhibition constants, k_i_. As can be seen from Figure 7a, this combination was identified quickly by the model with approximately optimal results being identified within 20 generations. Figures 7b, 7c and 7d show the dynamics of the model after this intervention is applied at forty years following forty years of the high risk lipid profile. Here we can see that smooth muscle cells, macrophages and foam cells and Th1-cell counts are all rapidly driven down by the intervention.

**Figure 7.**
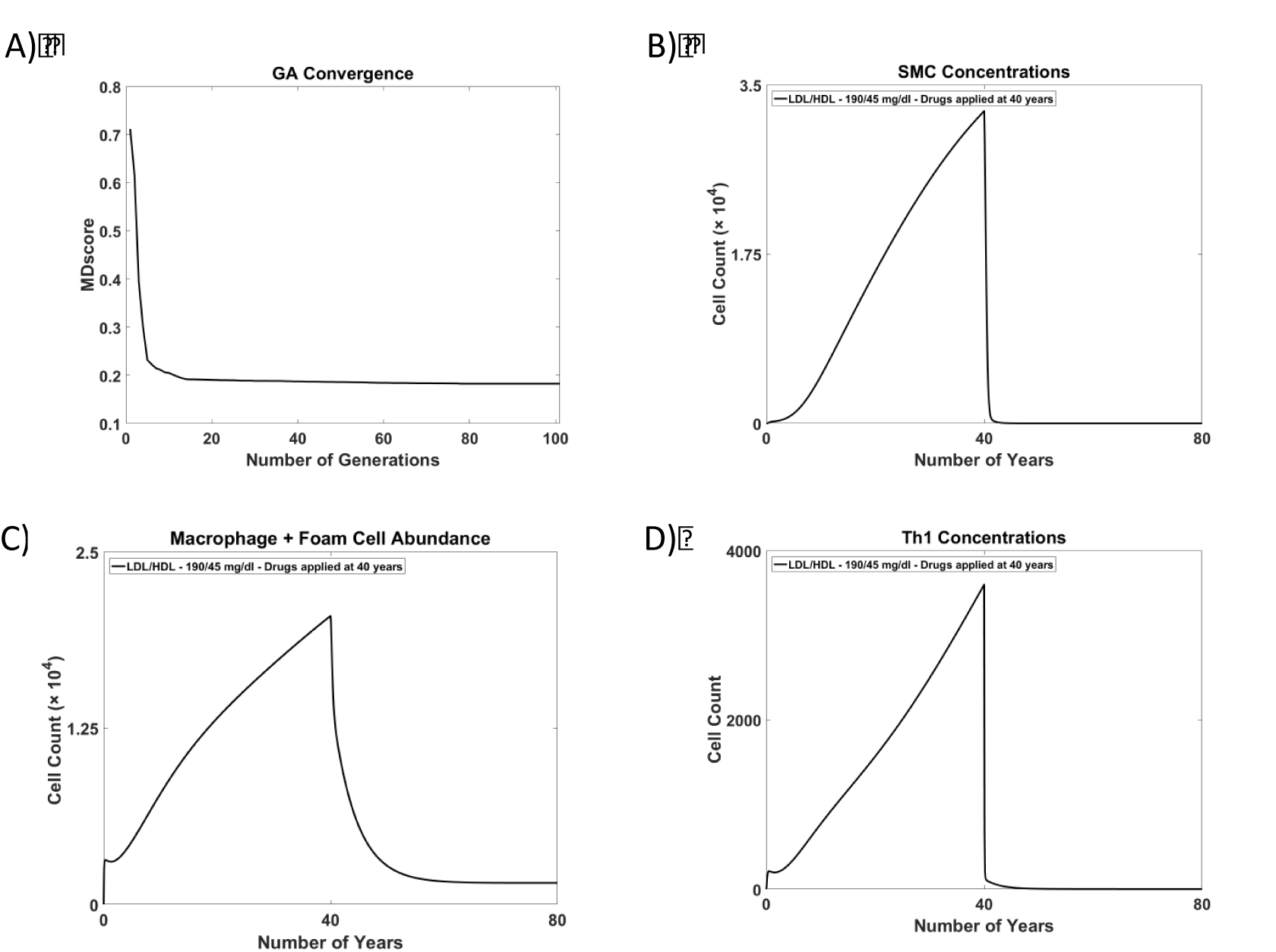
A) Convergence on an atheroprotective multi-drug intervention hypothesis. B-D) The impact of that intervention on key plaque components when applied after 40 years of a high risk LDL/HDL profile of 190/40mg/dl.

## 4. Discussion

Atherosclerotic plaques are highly challenging to study due to their location. *In vivo* study presents logistical and ethical challenges and there are few *in vitro* resources that can contribute to our understanding of plaque development. Whilst they are not a complete replacement for *in vivo* studies, computational studies have the potential to contribute to research in this area, and to yield non-*in vivo* resources that can improve our understanding of CVD.

CVD is a large burden on healthcare worldwide. Front line therapies for the primary and secondary prevention of atherosclerotic disease include smoking cessation, lipid management, blood pressure control, optimal control of diabetes and the use of antiplatelet agents. By far the most commonly used class of lipid lowering drugs is statins, which inhibit HMG CoA reductase. Ezetimibe, a cholesterol absorption inhibitor, may be used in patients who are statin intolerant or who do not achieve lipid targets on the highest maximally targeted dose of statin. A new, recently licenced class of drugs, proprotein convertase subtilisin/kexin type 9 (PCSK9) inhibitors, suppress degradation of LDLR by PCSK9 and are associated with a significant reduction in serum LDL concentration and in cardiovascular events. Emerging drugs include Apolipoprotein B antisense drugs that suppress translation of ApoB, a key component of LDL, and microsomal triglyceride transfer protein inhibitors that induce significant LDL reduction.

Here we have produced a predictive model of the dynamics of atherosclerosis, which we hope will serve as a resource for the cardiovascular research community that can be reused, refined and expanded in future. The model we have produced has the potential to contribute to therapy development through multiple avenues. Primarily, the model can be used to predict the consequences for the dynamics of atherosclerosis of interventions that target components of the pathways involved in the disease. This can be exploited in single drug development by identifying the components of the model that have the greatest impact on plaque development as potential drug targets or to multi-drug interventions that achieve similar goals through compound effects^41^. It is known that atherosclerosis is a comorbidity of diseases such as rheumatoid arthritis and depression^51^. By using proteomic data from studies of other diseases, this model can also be used to explore the role of atherosclerosis as a comorbidity. Furthermore, it can be used to explore the possible off-target impact of therapies for seemingly unrelated conditions, where the therapeutics are known to have targets within atherosclerosis associated pathways.

Although we often consider disease pathologies in isolation, atherosclerosis is part of a much larger network of interactions and we can use the model to explore the impact of interventions on the network of interactions that regulate atherosclerosis. For example, it would be possible to extend the model to include PCSK9 metabolism in order to explore the impact of PCSK9 inhibitors on plaque development or to include jak-stat signaling to explore the role of innate immunity on atherosclerosis progression.

The predictions of the model show broad agreement with observed clinical results. Because the model describes spatial effects and cellular function at extremely simple levels, it is unlikely to be able to recreate all clinical results exactly. Doing so would require a model of greater complexity across multiple length scales. However, the model presented here demonstrates order of magnitude agreement in almost all cases and shows the correct qualitative dose responses. We found it challenging to optimise the parameters so as to ensure a sufficiently large response to changes in lipoprotein profile for particular model components. As a result, particular components are systematically over-estimated for the low LDL profile and the difference between high and low LDL profiles, although large, is not as great as that observed clinically. In changing the lipid profile, we adjusted the concentrations of LDL and HDL in the model. This logically does not impact on the model components upstream of LDL and HDL. Hence, as observed in XXVIII and XXIX, we would expect to see no resulting change in chylomicron or triglyceride concentrations. To achieve this would require either generating VLDL and IDL values across patient risk profiles or incorporating greater feedback into the model.

A predictive model of this type has the potential to move the discussion around disease from an understanding of behavior of individual disease components (such as foam cell accumulation or smooth muscle cell recruitment) to an understanding of the dynamics of the network and of how the network as a whole transitions from healthy dynamics to disease dynamics.

As demonstrated, a model of this form can be used to develop therapeutic hypotheses. In principle, the model can be adapted to individuals or to patient subgroups by tuning the parameters of the interactions enabling it to contribute to programmes of personalized or stratified medicine. Parameterisations that are tailored to individuals could be identified by optimizing the model to patient or patient group time course data or from computational inference from single nucleotide polymorphism or genome data. Adapting the model to represent the disease dynamics of individual patients or patient subgroups would support the development of therapeutic hypotheses that are tailored to the patient or the patient subgroup.

The scale of the global CVD burden means that there is a pressing need to develop new pharmaceutical therapeutics in this area that both address clinical need and can sustain the pharmaceutical industry as intellectual property protection expires around current therapeutics. Multi-drug interventions of the type identified here have a vast untapped potential to contribute to future therapeutics in this way.

## Acknowledgements

We are indebted to Patricia Navarro for assistance with figure production.

## Footnotes

a http://www.escardio. http://org/static_file/Escardio/Press-media/press-releases/2013/EUcardiovascular-disease-statistics-2012.pdf

b https://www.ebi.ac.uk/biomodels-main/

c http://sbgn.github.io/sbgn/software_support

d http://sbml.org/SBML_Software_Guide

e https://www.nhlbi.nih.gov/health/resources/heart/heart-cholesterol-hbc-what-html

f https://www.mathsworks.com

g http://copasi.org/

h http://apps.cytoscape.org/apps/cy3sbml

i http://magnet.systemsbiology.net/software/Dizzy/

j http://web.newteditor.org/

k https://www.pathvisio.org/

l https://immersive-analytics.infotech.monash.edu/vanted/addons/sbgn-ed/

